# The birth-and-death evolution of cytochrome P450 genes in bees

**DOI:** 10.1101/2021.09.07.459278

**Authors:** Kathy Darragh, David R. Nelson, Santiago R. Ramírez

**Affiliations:** Department of Evolution and Ecology, University of California, Davis, CA, 95616; Department of Molecular Sciences, University of Tennessee, Memphis, TN, 38163

**Keywords:** cytochrome P450, birth-and-death evolution, gene family evolution, orchid bees

## Abstract

The birth-and-death model of multigene family evolution describes how families can expand by duplication and contract by gene deletion and formation of pseudogenes. The phylogenetic stability of a gene is thought to be related to the degree of functional importance. However, it is unclear how much evolution of a gene in a gene family is driven by adaptive versus neutral processes. The cytochrome P450s are one of the most diverse and well-studied multigene families, involved in both physiological and xenobiotic functions. Bees have a high toxin exposure due to their diet of nectar and pollen, as well as the resin-collecting behavior exhibited by some bees. Here, we describe the P450s of the orchid bee *Euglossa dilemma*. Orchid bees are a neotropical clade in which males form perfume bouquets used in courtship displays by collecting a diverse set of volatile compounds, resulting in high chemical compound exposure. We conducted phylogenetic and selection analyses across ten bee species encompassing three bee families. We do not find a relationship between the ecology of a bee species and its P450 repertoire. Our analyses reveal that P450 clades can be classified into stable and unstable clades, and that genes involved in xenobiotic metabolism are more likely to belong to unstable clades. Furthermore, we find that unstable clades are under more dynamic evolutionary pressures, with signals of adaptive evolution detected, suggesting that both gene duplication and positive selection driving sequence divergence have played a role in the diversification of bee P450s. Our works highlights the complexity of multigene family evolution which does not always follow generalized predictions.

**Significance statement:** Gene family evolution is characterized by deletion and duplication, but it is unclear if this is driven by adaptive or neutral evolutionary processes. To investigate the dynamics of gene family evolution we analyze the P450s of ten bee species, a well-studied gene family involved in both physiological and detoxification roles. We find that genes involved in detoxification are more likely to belong to unstable clades, exhibit dynamic evolutionary pressures, and show signals of adaptive evolution. However, we do not find evidence for a relationship between resin collection or sociality, and the P450 repertoire of a bee species. Our findings highlight the complexity of multigene family evolution which does not always follow generalized predictions.

## Introduction

Multigene families arise from the duplication of a common ancestral gene, resulting in groups of genes that share similar sequences, and often functions (Nei & Rooney 2005). To understand how these families evolve, early studies focused on concerted evolution in rRNA, whereby members of a gene family evolve together due to repeated unequal crossover events (Brown et al. 1972). More recently, an alternative model, the birth-and-death model, has been used to explain gene family evolution. In this model, genes evolve independently and expansion and contraction occurs through gene duplication, formation of pseudogenes, and gene deletion (Nei & Rooney 2005). The relative roles of neutral versus adaptive evolutionary forces within the framework of the birth-and-death model has proven more complex (Eirín-López et al. 2012).

Cytochrome P450s are one of the best-studied and most diverse multigene families (Nelson et al. 2013). They are enzymes which use molecular oxygen to change the structure of their substrates, a reaction which has an important role in both physiological endogenous processes, and also ecological and xenobiotic processes (Nelson et al. 2013). P450s are highly diverse and groups have been named based on sequence similarity, with four prominent groups described in insects (Nelson 1998; Tijet et al. 2001; Dermauw et al. 2020). Of these, the CYP3 group is the largest and most dynamic, exhibiting many lineage-specific duplications in insects, often linked to insecticide resistance and xenobiotic metabolism (Feyereisen 2006).

There appears to be a link between the function of P450s and their evolutionary dynamics. In vertebrates, genes involved in xenobiotic detoxification are more likely to be evolutionarily unstable, exhibiting duplications and deletions between species (Thomas 2007). In contrast, those involved in viability are more likely to be stable with a one-to-one orthology between species (Thomas 2007). This pattern has also been found in Drosophila where P450s associated with a role in development are duplicated less than those implicated in detoxification or have unknown functions (Drosophila 12 Genomes Consortium 2007).

Whilst the link between evolutionary instability in P450s and xenobiotic function has been shown, this does not test whether selection is acting on these P450s. The expansion of P450 subfamilies involved in xenobiotic functions is often assumed to be due to environmental adaptation driven by natural selection. However, even if an initial duplication is selected for, further duplication which lead to large subfamilies could simply be due to the self-sustaining process whereby a duplicated gene has twice the likelihood of duplicating again (Feyereisen 2011). In contrast to this neutral viewpoint, at a molecular level, correlation between amino acid replacement and the number of duplications found for a P450 lineage in the Drosophila phylogeny is inconsistent with stochastic models (Good et al. 2014). Furthermore, CYP expansions have been linked to environmental factors, such as specialized diets in Lepidoptera (Calla et al. 2017), implicating adaptive evolutionary forces in P450 expansions. However, the observed pattern of many groups with few genes and few groups with many genes (power-law distribution) does not require an adaptive explanation (Dermauw et al. 2020). Birth-and-death models of gene family evolution are sufficient to explain the pattern, not requiring any further explanation based on the ecology or life-history of the species (Sezutsu et al. 2013).

An interesting group for the study of P450s and detoxification is bees. Whilst providing important ecosystem services as pollinators, bees are exposed to a wide range of toxins, both natural and synthetic (Johnson 2015). In particular, bees differ from most pollinators in the fact that they are specialized on consuming pollen and nectar during all life-stages, diet sources with high levels of potentially toxic flavonoids, and have been called “flavonoid specialists” due to their consistent exposure to these compounds (Johnson et al. 2018). Exposure to flavonoids is thought to be higher in perennial eusocial bees due to the concentration of flavonoids when nectar is converted into honey and pollen into beebread for storage (Johnson et al. 2018). Furthermore, resin collection, associated with the evolution of sociality but also found in some solitary species, also increases flavonoid exposure (Bankova et al. 1983).

Understanding the molecular basis of toxin sensitivity is important to protect bees, especially from the negative effects of pesticides (Berenbaum & Johnson 2015). The sequencing of the honeybee genome (*Apis mellifera*) revealed a lower diversity of detoxification genes than expected when compared with other insect species (Claudianos et al. 2006; The Honeybee Genome Sequencing Consortium 2006). While the beetle *Tribolium castaneum* and the mosquito *Anopheles gambiae* have over 100 P450s, the honey bee *Apis mellifera* has less than 50 (Ranson et al. 2002; Claudianos et al. 2006; Oakeshott et al. 2010; Zhu et al. 2013). This deficit of P450s is not only exclusive to *Apis mellifera*, but is also the case in other bee species, including the solitary leafcutter bees *Megachile rotundata* and *Osmia bicornis* (Beadle et al. 2019; Hayward et al. 2019). Furthermore, this reduction is not equal across all P450s. Mitochondrial and CYP2 P450s have mostly been maintained, with the genes present from these groups linked to physiological roles in other insects such as hormone biosynthesis (Claudianos et al. 2006). In contrast, the CYP4s are greatly reduced, especially those members thought to play a role in xenobiotic detoxification. The CYP3s are also greatly reduced in diversity, but widespread duplication in the CYP6AS family within the CYP3s means that these make up a large proportion of total bee P450S (Claudianos et al. 2006; Berenbaum & Johnson 2015). This may explain why, in spite of these reductions, honey bees are not more sensitive to insecticides that other insects (Hardstone & Scott 2010).

One particular group of bees, the orchid bees, exhibit unique natural history and adaptation that presents additional challenges for detoxification. Male bees collect chemical compounds from different sources, such as orchid flowers and fungi, which they store in specialized leg pouches to concoct a ‘perfume’ bouquet (Dressler 1982). These bouquets are then released during courtship displays and are thought to play a key role in mate choice (Eltz et al. 2005; Pokorny et al. 2017). The chemical bouquets differ between species, but also between individuals of the same species and may provide information to the female regarding mate quality (Eltz et al. 1999; Zimmermann et al. 2009; Weber et al. 2016). Male orchid bees collect a wide range of chemical compounds, many of which are expected to be toxic to bees. Collection of these compounds, therefore, could act as an indicator of male quality, through their ability to handle and detoxify these compounds (Eltz et al. 1999; Arriaga-Osnaya et al. 2017). We may expect adaptation at the molecular level, perhaps in the form of an increased P450 repertoire, or increased expression levels, to be able to detoxify such a wide range of compounds.

Here, we study the molecular evolution of cytochrome P450s in bees and evaluate the hypothesis that orchid bees have an expanded P450 repertoire. We annotate P450s in the genome of *Euglossa dilemma* and combine these newly annotated P450s with those previously identified in nine other bee species: *Apis mellifera*, *Bombus terrestris*, *Dufourea novaeangliae*, *Eufriesea mexicana*, *Habropoda laboriosa*, *Lasioglossum albipes, Megachile rotundata, Melipona quadrifasciata,* and *Osmia bicornis.* These species are from three different families, and include *Eufresia mexicana*, another orchid bee. To determine the evolutionary history of these P450s we carry out phylogenetic analyses and classify clades as stable or unstable based on their history across these species. We measure evolutionary change and search for signals of adaptive evolution in these clades to identify the selection pressures acting on P450s in these bee species. We also investigate the patterns of correlation between the level of sociality of a bee species, whether or not it collects resin, and its P450 inventory. We aim to shed light both on the evolution of the P450 family and orchid bee biology in relation to detoxification.

## Results

### Phylogenetic analysis of bee P450s

To compile a dataset of bee P450s we firstly combined previously annotated P450s for nine species: *Apis mellifera, Bombus terrestris, Dufourea novaeangliae, Eufriesea mexicana, Habropoda laboriosa, Lasioglossum albipes, Megachile rotundata, Melipona quadrifasciata,* and *Osmia bicornis* (Hayward et al. 2019; Beadle et al. 2019; Kapheim et al. 2015; Johnson et al. 2018) (Figure 1). To this set of P450s we then added 41 P450s which we identified and annotated in *Euglossa dilemma* with complete protein domains. We also identified 4 P450s with incomplete protein domains which may represent pseudogenes or may be due to poor assembly, sequencing errors, or sequencing gaps. We do not include these for the following analyses (incomplete_genes.fa).

**Figure 1:**
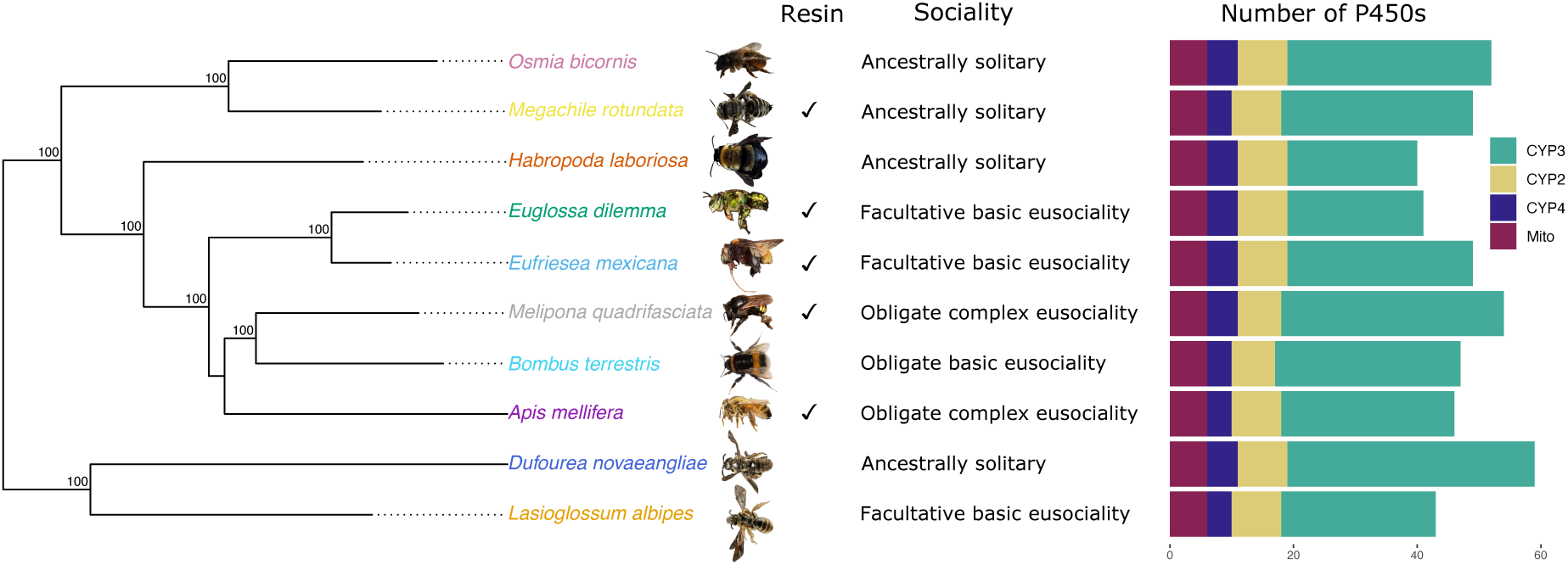
Phylogeny of cytochrome P450s in bees. The phylogeny was constructed in IQ - TREE (model *GTR+F+R4*) using five nuclear genes at 10508 nucleotide sites. Bootstrap values (n=1000) are illustrated. For each species its resin collecting behavior, level of sociality, and number of P450s in each family is shown. *Melipona quadrifasciata* photo courtesy of Lucas Rubio, *Osmia bicornis* photo courtesy of Amelia Bassiti.

To determine orthology of P450s we constructed a phylogeny combining the *Eg. dilemma* P450s from those previously identified in nine other bee species. The P450s group into the four expected clusters in insects, CYP2, CYP3, CYP4 and mitochondrial (Figure 2, Supplementary Figure 1). As expected for bees, the CYP4s are reduced across all species, and the CYP6AS family is expanded. In some clades the phylogeny of P450s does not reflect the species’ phylogeny. This is likely due to errors in phylogenetic reconstruction in these particular P450 clades rather than complicated patterns of duplicated and deletion between species. No members of the CYP6AS clade containing CYP6AS10 from *B. terrestris* and *A. mellifera* are found in either *Eg. dilemma* or *Ef. mexicana*, suggesting this is a loss in orchid bees. Overall, *Eg. dilemma* has a comparable number and distribution of P450s as other previously studied bee species (Figure 1, Supplementary Figure 1).

**Figure 2:**
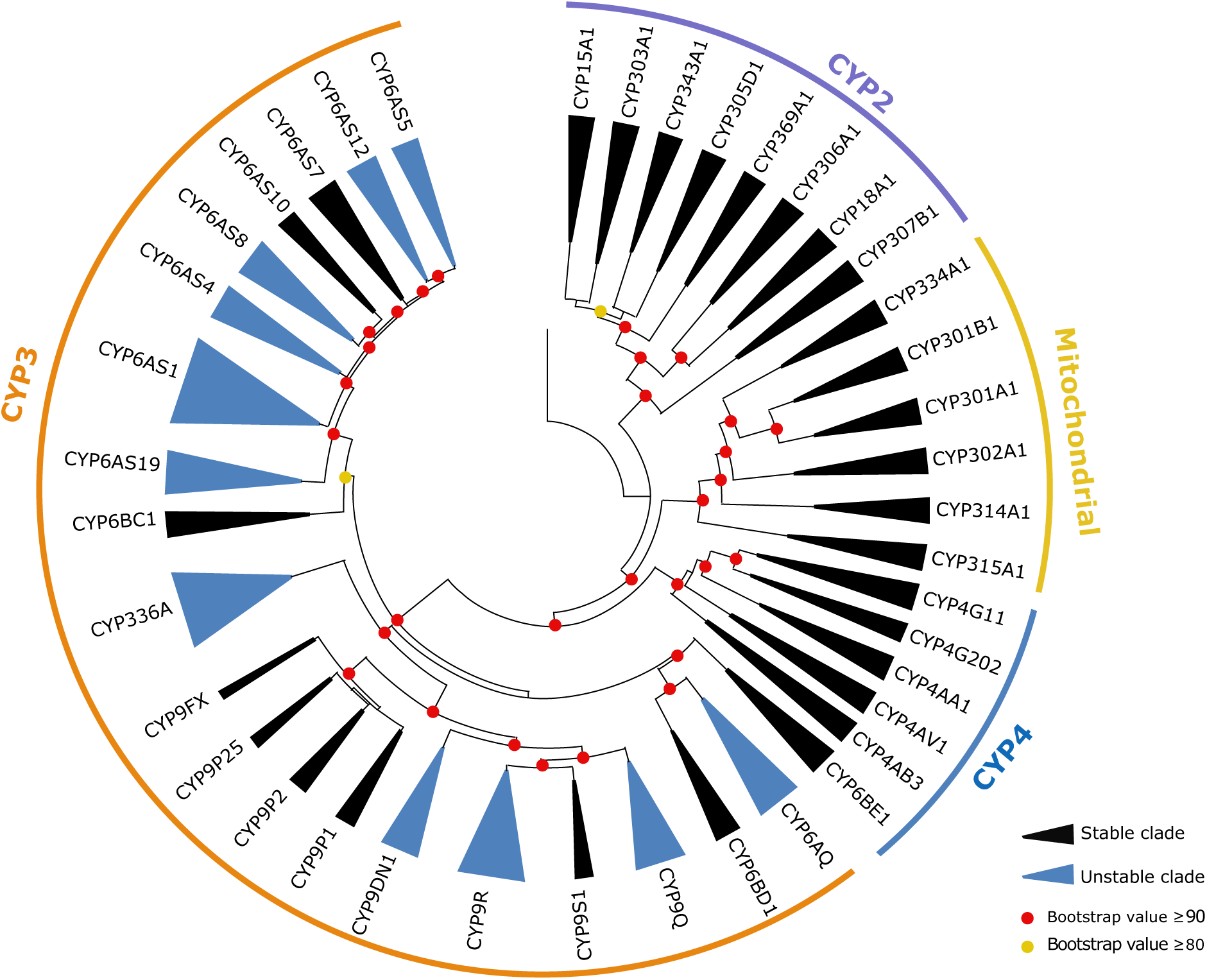
Phylogeny of cytochrome P450s in bees. The phylogeny was constructed in IQ -TREE (model JTT+F+R10) using 481 amino acid sequences across ten bee species belonging to three different families. Clades are collapsed to show 40 clades. Bootstrap values (n=1000) are illustrated.

### Is the number of P450s in the genome associated with species’ ecology?

We found that total number of P450s in the genome of each species ranged from 40 in *H. laboriosa* to 59 in *D. novaeangliae,* with most of this variation from the number of CYP3s found in each species (Figure 1). We investigated two aspects of species’ ecology: resin collection and level of sociality. We tested for relationships between P450 number and ecology while accounting for phylogeny. We did not find a relationship between sociality and the number of P450s (Supplementary Figure 2A), the number of CYP3s (Supplementary Figure 2B), or the number of CYP6AS genes (Supplementary Figure 2C). Furthermore, we did not find a difference between bees that exhibit substantial resin collection and those which do not in the number of P450s (Supplementary Figure 3A), the number of CYP3s (Supplementary Figure 3B), or the number of CYP6AS genes (Supplementary Figure 3C).

### Evolutionary history of P450s in bees

To investigate the dynamics of gene family evolution and phylogenetic instability there are two main approaches: the reconciliation of the gene tree with the species tree to detect duplication and loss events, and the use of birth-death models fitted to a species tree (Yohe et al. 2019).

We used MiPhy to implements the first approach of species tree gene tree reconciliation to identify clades and estimate phylogenetic instability scores for each clade. Without the “merge singletons” option, MiPhy divided our data into 42 clades including two clades which only contained a sequence from *Habropoda laboriosa.* For the final analysis, we used the option “merge singletons” to divide our data into 40 clades ranging in their instability score from -0.1 to 25.05 (Supplementary Figure 4). The instability scores of the clades fell into two broad groups. Clades with low instability scores (-0.1 to 3.61) were classified as stable, and those with higher instability scores (9.12-25.05) were classified as unstable. In total, 27.5% (11/40) clades were classified as stable. The distribution of stable and unstable clades was not equal amongst P450 families, with 52% of CYP3 clades found to be unstable, in comparison with only 0% of CYP2, CYP4 and mitochondrial P450 clades (Figure 2). CYP3s have a higher instability score than the other CYP groups (Supplementary Figure 5).

To apply the second approach to our dataset we use Computational Analysis of gene Family Evolution (CAFÉ), a program which uses birth-death to model gene gain and loss across a species tree, using the clades previously identified by the MiPhy analysis (Mendes et al. 2020). Clades which are fast evolving are identified by comparing models in which all clades evolve at the same rate with models in which different clades vary in their evolutionary rate. Using this approach, we identified one CYP6AS clade, CYP6AS1, as an outlier when compared to all other families, with respect to its evolutionary rate of family size change (p=0.029). This was the same clade which had the highest instability in the MiPhy analysis (Supplementary Figure 4). As expected, this family also had the strongest assignment to the gamma category of the highest median lambda (rate of evolutionary change), (posterior probability=1.0). Furthermore, whilst not detected as outliers relative to the entire dataset, seven additional clades were also inferred to be rapidly evolving in size, as indicated by their significant placement in the same fast evolving gamma category (Supplementary Table 1; posterior probability > 0.95). All of these were classed as unstable by the MiPhy analysis. Whilst all of the clades identified by CAFE were identified as unstable in the MiPh analysis, this was not true in reverse, probably as CAFE uses gene counts and does not consider the gene tree, making it unable to distinguish between independent gene duplication events and inheritance of paralogs (Curran et al. 2018).

The exact pattern of duplications and deletions is unclear, but certainly some have happened on a more recent timescale. For example, CYP9DN1 is not found in *A. mellifera*, *B. terrestris*, *D. novaeangliae*, or *Ef. mexicana*, suggesting this deletion has occurred at least twice as *D. novaeangliae* is not closely related to the other species. Of the 11 unstable clades, three belong to this category of requiring at least two deletion events to explain the phylogenetic pattern. Other clades could be explained by one deeper internal deletion event, but more species would be needed to determine this. For example, the clade containing *Apis mellifera* CYP6AS10 is missing in both *Ef. mexicana* and *Eg. dilemma*, which could be explained by a deletion in all orchid bees, or two separate deletion events, with further sampling necessary to distinguish the two. Duplication events are also a mix of lineage-specific and potentially deeper internal duplications. The genes CYP6AS131-135 and CYP6AS91-95 are expanded specifically in *O. bicornis* and *D. novaeangliae*, respectively, suggesting more recent duplication events. In contrast, CYP336 shows a pattern consistent with duplications deeper in the lineage leading to *O. bicornis* and *M. rotundata* (Supplementary figure 1).

### Do unstable clades of P450s exhibit increased evolutionary change?

The clades were found to differ in their phylogenetic instability phylogenetic instability and rate of size evolution. To further investigate the evolutionary dynamics of these clades we compared various evolutionary measures. Branch length can be used as a measure of evolutionary change in a phylogenetic tree. We found a correlation between instability and branch length, when gene number is corrected for, suggesting higher rates of evolution than for genes found in stable clades (Fig 3). Unstable clades also have a higher cumulative patristic distance, which includes the length of internal branches also (Supplementary Figure 6). One clade, CYP9FX, did not follow this trend, with a low instability score but high CBL and cumulative patristic distance. This is the smallest clade in our analyses with only three genes from *Dufourea novaeangliae* and *Lasioglossum albipes*.

**Figure 3.**
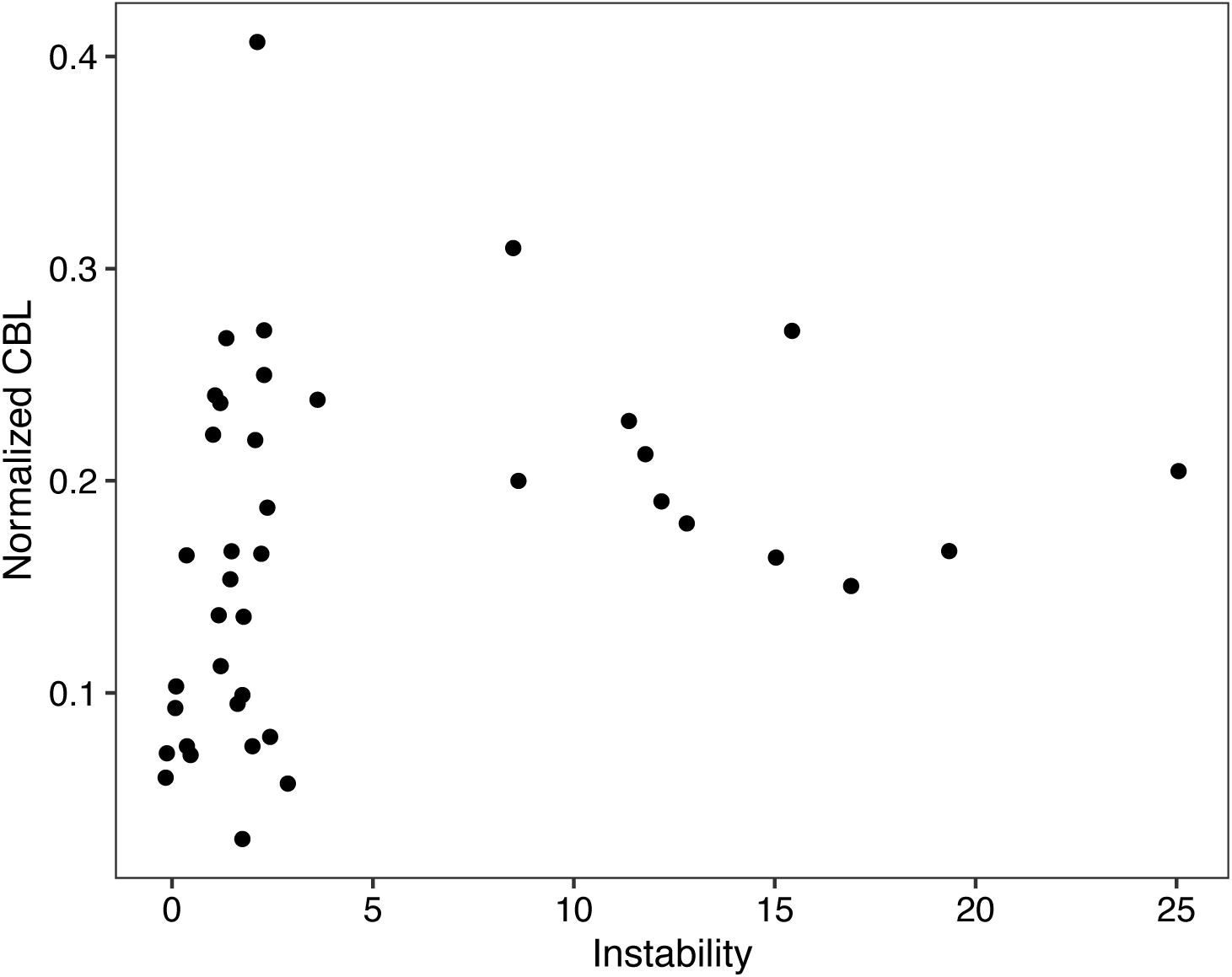
Relationship between CBL and the instability of a clade. When CBL is normalized for gene number in a clade, there is a significant correlation between clade CBL and clade instability (Spearman’s Correlation, rho=0.401, p= 0.0103). CBL, cumulative branch length.

### Are there signals of adaptive evolution in bee P450s?

#### Branch-specific models

Although branch length can give an indication of the amount of divergence between two P450s, it does not provide information on the predominant type of selection that shaped their evolution. To explore both the amount and the type of evolutionary change happening in stable and unstable clades, we compared the rate of nonsynonymous to synonymous mutations (dN/dS) in different lineages. The simplest analysis carried out was a “one-ratio” model for each clade. In these models, each branch in a clade is assumed to have the same dN/dS value. All values obtained were below one when all members of a clade are forced to have the same dN/dS value, suggesting that in general purifying selection is acting. To test if unstable and stable clades are under different selection pressures, we compared their “one-ratio” dN/dS values. We found that all clades are under purifying selection with dN/dS values less than one. However, we detected a correlation between clade instability and dN/dS ratio, with more unstable clades having a higher dN/dS ratio (Fig. 4A). Furthermore, we found a correlation between the cumulative branch length of a clade, the amount of evolutionary change, and the dN/dS value of the clade (Fig. 4B).

**Figure 4.**
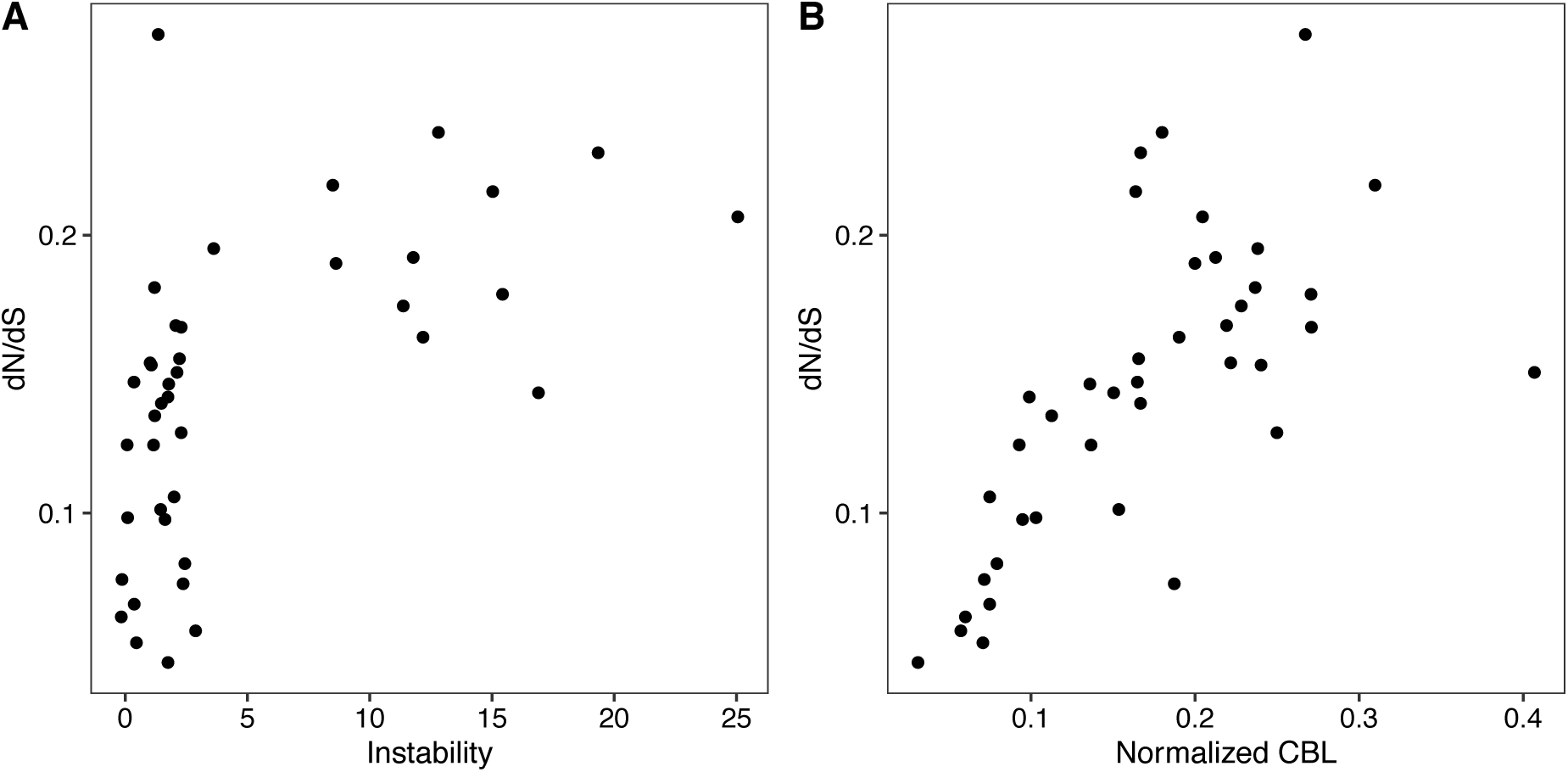
Relationship between the dN/dS of a clade, clade instability, and normalized CBL. A) Correlation between the instability of a clade and its dN/dS ratio in “one-ratio” models (Spearman’s Correlation, rho=0.567, p= 0.00014). B) Correlation between Normalized CBL (average branch length per gene in a clade) and dN/dS of the clade in a “one-ratio” model (Spearman’s Correlation, rho=0.754, p= 1.889e-08). CBL, cumulative branch length.

The “one-ratio” model is restrictive, only allowing a single dN/dS value per clade. To investigate which clades are under more dynamic selection pressures than the simple “one-ratio” model, we compared these models to models under which each lineage can vary in its dN/dS ratio, also called “free-ratio” models (Supplementary Table 2). As for the “one-ratio” models above, these “free-ratio” models were carried out by modelling each clade individually. The evolutionary history of unstable clades is more dynamic than stable clades. The more complex model has a better fit for more of the unstable clades (100%, 11/11) than the stable clades (69%, 20/29) (Fisher’s exact test, p=0.0433)).

As well as testing selection pressures at a clade level we tested each branch in each clade individually for evidence of selection. We compared “one-ratio” models for each clade with models in which only one branch is allowed to vary in its dN/dS ratio (“two-ratio” models). In these models we can ask whether the branch of interest has a dN/dS value significantly different from the “background” dN/dS of the clade. Again, we found the same pattern in the resulting dN/dS values. Of all the branches with a dN/dS ratio significantly different from the background of the clade, dN/dS was higher for branches from unstable than stable clades (Supplementary Figure 7). Furthermore, all branches with a significant dN/dS greater than 1.1 (excluding branches where dS<0.01 and dS>2), indicating positive selection, were from the CYP3 clade. Of these, 70% (7/10) were from unstable CYP3 clades, and 70% (7/10) were from the CYP6AS family (Table 1). The three genes which were not found in unstable clades were all in the CYP6AS7 clade. Full list of tests can be found in Supplementary data 2.

**Table 1:** Branches with dN/dS greater than one and significantly different from background dN/dS of the clade in the “two-ratio” tests (adjusted p values <0.05). Branches with dS<0.01 and dS>2 have been removed.

| Name | dN/dS | CYP<br>clade | CYP family | Genus |
| --- | --- | --- | --- | --- |
| GB40288 | 2.316 | CYP3 | CYP6AS1 | *Apis mellifera* |
| LALB19295RA_B | 1.822 | CYP3 | CYP6AS200 | *Lasioglossum*<br>*albipes* |
| MQUA24239RA | 1.597 | CYP3 | CYP336A42 | *Melipona*<br>*quadrifasciata* |
| Internal branch | 1.597 | CYP3 | CYP336 |  |
| MH500637 | 1.322 | CYP3 | CYP6AS127 | *Osmia bicornis* |
| NW_015373885.1C | 1.268 | CYP3 | CYP9FW6 | *Dufourea*<br>*novaeangliae* |
| Internal branch | 1.243 | CYP3 | CYP6AS7 |  |
| MH500655 | 1.224 | CYP3 | CYP6AS131 | *Osmia bicornis* |
| MH500653 | 1.206 | CYP3 | CYP6AS132 | *Osmia bicornis* |
| XM_017902249.1 | 1.137 | CYP3 | CYP6AS87 | *Eufriesea*<br>*mexicana* |

#### Site-specific models

Branch-specific models assumes a consistent dN/dS ratio across all sites in a given gene. We also carried out site-specific tests which allow dN/dS to vary among sites. We found evidence of positively selected sites in a quarter (10/40) of all clades in the M7 and M8 model, including clades from CYP2, CYP4 and CYP3 (Supplementary Table 3). For four of these clades, CYP6AS1, CYP6AS7, CYP6AS8 and CYP336, positively selected branches were previously identified in the branch-specific test comparing “one-ratio” and “two-ratio” models. CYP6AS1 and CYP6336 are the two clades with highest instability as identified by MiPhy and CYP6AS1 was also identified as the sole outlier family in terms of evolutionary rate by CAFE. We identified positively selected sites at 90% posterior probability for eight of these clades: CYP306A1, CYP305D1 CYP4G202, CYP336, CYP6BD1, CYP6AS1, CYP6AS7, and CYP6AS8. We found no evidence of positively selected sites in the M1a-M2a comparison.

To determine where on the protein these sites were located, we took advantage of the AlphaFold2 algorithm (Jumper et al. 2021; Mirdita et al. 2021) to create protein structural models for a representative sequence in each clade. In general, the residues are found on surface regions of the proteins. The resides identified in CYP6AS1, CYP6AS7, CYP6AS8 and CYP6BD1 are all found in the same surface region of the protein (Supplementary Figure 8). The residues in CYP6AS1 and CYP6AS7 are both found between the F’ and F helices, and those for CYP6AS8 and CYP6BD1 are both found between the G and G’ helices. This region is not close to the active site, and may be involved in protein-protein interactions (Sevrioukova et al. 1999). The residues for the other four P450s are found in other regions of the structure (Supplementary Figure 9). The positively selected site identified in CYP306A1 is found at the C-terminal of the protein. Two residues were identified in CYP305D1, one at the end of the F helix and the other on a surface loop between the I and H helices. Two were also detected for CYP4G202, one in the turn between beta 1 and beta 2 near the N-terminal, and the other on a large surface loop. Finally, the residue in CYP336 is located on a non-helical, non-beta surface area of the protein. None of these residues are part of the active site of the protein.

## Discussion

Whilst the role of duplication and deletion in the evolution of multigene families is well-established, controversy remains surrounding whether this evolution is primarily adaptive or neutral. In this study, we demonstrate that genes known to be involved in xenobiotic metabolism (CYP3 family) are more likely to be unstable in the form of more gene duplication and deletion. Furthermore, we find that not only are these genes more likely to be unstable but also under more dynamic evolutionary pressures, and exhibit signals of adaptive evolution. This suggests that both gene duplication and positive selection driving sequence divergence contribute to the diversification of P450s.We do not find evidence for a correlation between the P450 repertoire of a bee species and its biology. Furthermore, our hypothesis that orchid bees would have an expansion of P450s due to their perfume collection behavior was also not supported.

The patterns we identify here are very similar to those previously identified in *Drosophila*. P450s with developmental functions are more likely to be found in stable clades, and also more likely to be under purifying selection (Drosophila 12 Genomes Consortium 2007; Good et al. 2014). These patterns, however, are not restricted to P450s, and have also been identified in the fatty acyl-coA synthase multigene family in Drosophila, which are involved in both essential physiological and chemical communication functions (Finet et al. 2019). While no single evolutionary model will be able to explain multigene family evolution, we can understand some common underlying principles using the framework of the birth-and-death model (Eirín-López et al. 2012).

Another common pattern found across multigene families is lineage-specific expansions, which are often associated with the evolution of novelty (Lespinet et al. 2002). In the CYP3 clade of P450s in bees, we find extensive evidence of duplications, particularly in the CYP6AS subfamily. It has been suggested that this expansion is related to specialization on a plant-based diet and the ability to metabolize flavonoids, in comparison to solitary carnivorous ancestors such as solitary wasps (Johnson et al. 2018). As previously described by Johnson and colleagues (Johnson et al. 2018), we find that not only are CYP6AS subfamilies expanded by duplication, but also show evidence of positive selection, with the majority of positively selected genes detected belonging to this family. CYP6AS3 and CYP6AS4 from *Apis mellifera* were found to be under positive selection by Johnson and colleagues. Here we find CYP6AS1 to be under positive selection in the branch-specific models and evidence for positively selected sites in the clade containing CYP6AS1 and CYP6AS3 in the site-specific models. CYP6AS1, CYP6AS3, CYP6AS4, and CYP6AS10, have all been linked to quercetin metabolism, a plant flavonoid found in honey (Mao et al. 2009). Other CYP6AS genes, as well as CYP9R1, CYP9P1, CYP9S1, and CYP9Q family genes are upregulated in response to quercetin treatment in *Apis mellifera* (Mao et al. 2017). Furthermore, CYP9Q4 in *Bombus* and CYP9Q3 in *Apis* metabolize the neonicotinoid thiacloprid (Manjon et al. 2018). All of these genes known to be involved in xenobiotic metabolism were identified in our study as members of unstable clades, again linking instability and gene duplication with xenobiotic function.

In contrast to these genes with known xenobiotic function, we find that genes predicted to have physiological roles are mostly found in stable clades under purifying selection. The mitochondrial clade genes CYP302A1, -314A1, AND -315A1, which are orthologs of the *Drosophila melanogaster* Halloween genes, are thought to be involved in ecdysteroid synthesis and are all found in stable clades in our analysis (Gilbert 2004; Rewitz et al. 2007; Feyereisen 2011). From the CYP2 clade, CYP18A1, thought to be involved in ecdysteroid inactivation, CYP15A1, a predicted juvenile hormone epoxidase, and CYP306A1, a CYP2 member orthologous to a *D. melanogaster* ecdysteroid 25-hydroxylase, are also found in stable clades (Helvig et al. 2004; Niwa et al. 2004; Claudianos et al. 2006). However, CYP306A1, despite being a stable clade is also one of the clades with evidence for a positively selected residue. Multiple other stable clades also showed evidence of positive selection in site-specific models (CYP305D1, CYP4G202, CYP6AS7 and CYP6BD1). Furthermore, the dichotomy of physiological and xenobiotic genes is not so simple, as both can be found in phylogenetic proximity (Dermauw et al. 2020). Furthermore, members of the same subfamily may have divergent roles. For example, while the CYP6AS subfamily is known to play a role in xenobiotic detoxification, CYP6AS8 and CYP6AS11 of *Apis mellifera* are expressed in mandibular glands, potentially involved in fatty acid signal synthesis (Wu et al. 2017; Dermauw et al. 2020). This highlights the complexity of multigene family evolution which does not always follow generalized predictions.

If P450 function is linked to their evolutionary dynamics, we might expect species with different biology to differ in their P450s (Rane et al. 2019). Specifically, we hypothesized that *Euglossa dilemma* would exhibit an expanded repertoire of P450s when compared with other bees, due to its perfume-collecting behavior increasing exposure to different chemical compounds. In contrast, we found that *Eg. Dilemma* has a comparable set of P450s similar to other bees, with no significant expansions detected. This does not rule out an increased capacity for xenobiotic metabolism, which would need to be tested experimentally. Specialization could also occur at the sequence level rather than the number of genes; however, we did not find evidence for positive selection in *Eg. Dilemma* P450s. One alternative is that *Eg. Dilemma* exhibits increased gene diversity at another step in the detoxification pathway. In honeybees, not only are P450s reduced in comparison to other insects, but also glutathione-S-transferases (GSTs) and carboxyl/cholinesterases (CCEs) (Claudianos et al. 2006; Berenbaum & Johnson 2015). Both GSTs and CCEs are known to be involved in xenobiotic metabolism in insects, thus meriting further investigation in *Eg. Dilemma* (Li et al. 2007). Another possibility is that orchid bees do not demonstrate higher abilities to metabolize xenobiotic compounds, but instead reduce the amount of compound that can enter the body, for example by decreasing cuticle penetration, as reported in *Anopheles gambiae* (Balabanidou et al. 2016).

We also might expect a reduction in P450 repertoire in species which have a potentially lower xenobiotic exposure due to a specialist lifestyle (Rane et al. 2019). *Drosophila sechellia*, a specialist island species only found within a narrow ecological niche exhibits an increased loss of P450 genes compared to other *Drosophila* species (Good et al. 2014). In our dataset, the two most specialized species are *D. novaeangliae*, a specialist, with only one known pollen source (Eickwort et al. 1986), and *H. laboriosa*, with only a few known pollen sources (Pascarella 2007). Interestingly, *D. novaeangliae* has the largest P450 repertoire of all species in our analysis and *H. laboriosa* has the fewest P450s. We therefore do not find a strong link between specialization in diet and P450 repertoire in our bee dataset. More generally, we did not find a relationship between P450s and the biology of different bee species, neither at the total number level or subfamily level. This is in contrast to the previously identified trend that degree of sociality correlates with CYP6AS subfamily size, and that resin-collecting bees have more CYP6AS members (Johnson et al. 2018). This is likely because of the inclusion of *Osmia bicornis* in this study, a solitary bee which does not collect resin and yet has a large P450 repertoire, and the inclusion of phylogeny in these analyses. Future studies with more species, in particular species such as specialists, will allow us to untangle the relationship between species’ biology and P450s.

The P450 family in bees can provide us with insights into the evolution of multigene families. In addition to this evolutionary perspective, understanding detoxification in bees is crucial to native bee conservation and the management of agricultural pollination in the face of widespread pesticide use. Bees provide an important ecosystem service through pollination, both of wild plants and economically important crop species (Klein et al. 2007; Vanbergen & Initiative 2013). Whilst most insecticide assessments are carried out in *Apis mellifera*, they do not always reflect the sensitivity of native bees, with variation between species (Arena & Sgolastra 2014; Franklin & Raine 2019). Toxicity assays require large numbers of bees, difficult to achieve with many native bee species. Understanding the genetic mechanisms underlying insecticide sensitivity may allow us to make predictions based on P450 repertoires and sequences in different species (López-Osorio & Wurm 2020). A recent example of this comes from *Megachile rotundata*, a species which exhibits increased sensitivity to neonicotinoids, and also lacks the P450 enzymes known to be involved in neonicotinoid metabolism in other species (Hayward et al. 2019). A combination of comparative genomic studies, alongside toxicity assays and molecular and functional studies of metabolism and insecticide resistance will allow us to extend our understanding to other bee species.

## Materials and Methods

### Gene family annotation

First, we gathered previously annotated P450s for nine species: *Apis mellifera*, *Bombus terrestris*, *Dufourea novaeangliae*, *Eufriesea mexicana*, *Habropoda laboriosa*, *Lasioglossum albipes, Megachile rotundata, Melipona quadrifasciata,* and *Osmia bicornis* (Hayward et al. 2019; Beadle et al. 2019; Kapheim et al. 2015; Johnson et al. 2018). We combined all P450s described in previous publications and aligned them using MAFFT v7.453 (-maxiterate 1000, using L-INS-I algorithm) (Katoh et al. 2002, 2005; Katoh & Standley 2013). We then used the *bio3d* package in R to compare the sequences, removing duplicates to curate a final dataset (Grant et al. 2021), We searched either the National Center for Biotechnology Information (NCBI) or the Hymenoptera Genome Database to provide accession numbers for the included P450s. (Elsik et al. 2018; NCBI Resource Coordinators 2018). For genes not currently included in an annotation but previously manually annotated, we included the scaffold accession number where the sequence can be found (Supplementary data 1). We checked each sequence for a complete protein domain using the NCBI conserved domain search (Marchler-Bauer et al. 2015). In some cases the sequence from the database contained an incomplete domain which had been manually edited in the dataset provided by Johnson and colleagues (Johnson et al. 2018), and in these cases we used the manually curated sequence and noted this (Supplementary data 1).

We then annotated the P450 gene family in the *Euglossa dilemma* genome (Brand et al. 2017) using all previously annotated *Apis mellifera* protein sequences as a reference (Elsik et al. 2018). Firstly, we identified scaffolds in the *Eg. Dilemma* genome containing P450s using TBLASTN (Altschul et al. 1990), and then annotated proteins on these scaffolds using exonerate (Slater & Birney 2005). Gene models were extracted, translated to amino acid sequences and checked by comparison to *Apis mellifera* proteins. Gene models were manually curated using IGV, checking for splice sites and start and stop codons (Robinson et al. 2011). To check for P450s missed by the initial search, the predicted *Eg. Dilemma* P450s, as well as previously described *Ef. mexicana* P450s, were used as a query for the initial step of TBLASTN followed by exonerate predictions (Altschul et al. 1990; Slater & Birney 2005). As a further check we used already published RNA-seq data of ovaries and brains of *Eg. Dilemma* (Bioproject PRJNA523381) to improve the annotations (Saleh & Ramírez 2019). We trimmed the reads using TrimGalore! (Martin 2011). We then mapped the genes (2pass) to the *Eg. dilemma* genome (Brand et al. 2017) with the manually edited annotation (Edil_v1.0_revised.gff) using STAR (Dobin et al. 2013). We used StringTie (Pertea et al. 2016) to assemble and merge the transcripts to form an annotation which we compared to the edited annotation in IGV (Robinson et al. 2011). To check that the predicted P450s contain complete functional protein domains we used the NCBI conserved domain search (Marchler-Bauer et al. 2015). One gene, Edil_14289, is found at the end of a scaffold in a poorly sequenced area. The exon from the unsequenced area is found at the end of another scaffold and we combined these to form a complete gene in our analyses. All P450s identified were included in a revised version of the current *Eg. dilemma* annotation (Edil_v1.0_revised.gff) and included in Supplementary data 1.

### Phylogenetic analyses

All 481 bee P450 sequences were combined with CYP51 from *Mus musculus* as an outgroup. The sequences were aligned using MAFFT v7.453 (-maxiterate 1000, using L-INS-I algorithm) (Katoh et al. 2002, 2005; Katoh & Standley 2013). Phylogenetic trees were constructed using IQ-TREE with the ModelFinder function to determine the best-fit model (Nguyen et al. 2015; Kalyaanamoorthy et al. 2017; Hoang et al. 2018). The Newick phylogeny was plotted using MEGA X, and the packages *ape*, *evobiR*, and *geiger* in R version 3.6.2 (Pennell et al. 2014; Blackmon & Adams 2015; Kumar et al. 2018; Paradis & Schliep 2018; R Core Team 2019; Stecher et al. 2020).

To test for correlation between P450 repertoire and bee biology we carried out phylogenetic comparative analyses. The dataset used to produce the species-level phylogeny was composed of five nuclear genes: wingless, arginine kinase, opsin, Nak, Ef1α (F2 copy) (Danforth et al. 2011). We identified copies of the five genes in *Apis mellifera* from NCBI (NCBI Resource Coordinators 2018). We then used TBLASTN to search the nucleotide datasets on NCBI for the sequences in *Bombus terrestris*, *Dufourea novaeangliae*, *Eufriesea mexicana*, *Habropoda laboriosa, Megachile rotundata,* and *Osmia bicornis* (Altschul et al. 1990). For *Euglossa dilemma*, *Lasioglossum albipes* and *Melipona quadrifasciata*, we used TBLASTN and exonerate to find and annotate the proteins in the genome (Altschul et al. 1990; Slater & Birney 2005; Kapheim et al. 2015; Kocher et al. 2013; Brand et al. 2017). The sequences were aligned using MAFFT v7.453 (-maxiterate 1000, using L-INS-I algorithm) (Katoh et al. 2002, 2005; Katoh & Standley 2013). Alignments were trimmed using trimAl (- gt 0.6, sites only included where sequence from 6/10 bee species present) (Capella-Gutierrez et al. 2009), and trimmed alignments were then concatenated. We constructed phylogenetic trees constructed using IQ-TREE with the ModelFinder function to determine the best-fit model (Nguyen et al. 2015; Kalyaanamoorthy et al. 2017; Hoang et al. 2018), and constrained the relationship of Corbiculate bees according to a previous phylogeny (Romiguier et al. 2016). We rooted the tree using the midpoint.root function in the *phytools* package in R (Revell 2012).

Bees were determined as resin collecting or non-resin collecting following Johnson and colleagues (Johnson et al. 2018). We designated level of sociality as described previously (1= ancestrally solitary, 2=facultative basic eusociality, 3=obligate basic sociality, 4=obligate complex eusociality) (Kapheim et al. 2015). We tested for correlation between number of P450s, CYP3s, or CYP6AS genes and the level of sociality by using a Spearman’s rank correlation on phylogenetically independent contrasts (PICs). We calculated PICs using the function pic in the *ape* package (Paradis & Schliep 2018). We tested for a relationship between resin collection and P450 repertoire, using phylogenetic ANOVA, with the function aov.phylo from the package *geiger* (Pennell et al. 2014).

### Clade stability and branch length analyses

We used MiPhy to identify clades and assess their stability (Curran et al. 2018). MiPhy requires a rooted tree as an input, however, our root did not belong to any of the species present in the species tree. Therefore, we removed the *Mus musculus* CYP15 from the dataset, aligned the sequences using MAFFT v7.453 (-maxiterate 1000, using L-INS-I algorithm) (Katoh et al. 2002, 2005; Katoh & Standley 2013), and constructed a phylogenetic tree using IQ-TREE with the ModelFinder function to determine the best-fit model (Nguyen et al. 2015; Kalyaanamoorthy et al. 2017; Hoang et al. 2018). We rooted the tree using the midpoint.root function in the *phytools* package in R (Revell 2012). We used default parameter settings along with the option “merge singletons” to ensure that all clades had more than one gene. Clades were named using nomenclature from *A. mellifera*, for the CYP6AS subfamily which contained multiple clades, we named the clades using the lowest numbered group member from *A. mellifera*.

To compare instability between the four CYP groups, we used a Welch one-way ANOVA test, followed by a post-hoc Games-Howell test using the package *rstatix* (Kassambara 2021). These tests do not assume homogeneity of variance.

We then applied CAFE to our dataset to analyze the evolutionary rates of different clades. This program uses a birth-death process to model gene gain and loss across a species tree to identify fast-evolving clades. The input files required are a species tree and gene counts for each species in each clade, in this case the clades identified by MiPhy. We used the python script provided in the CAFE tutorial with the program r8s to convert the species tree constructed as described above into an ultrametric tree, as required for CAFE input (Sanderson 2003). We calibrated the tree using a previously published time-calibrated phylogeny (Cardinal et al. 2018). We found that three gamma rate categories (k=3) was the best fit for our data and showed convergence between runs.

For each clade we determined the cumulative branch length (CBL) by adding the terminal branches for each gene within a clade, obtained from the IQ-TREE output. As each clade has a different number of genes, we normalized the CBL by dividing the CBL by the number of genes in the clade, following (Finet et al. 2019). We also calculated the cumulative patristic distance per clade by summing all branch lengths, including internal branches within the clade. Again, we normalized the cumulative patristic distance by dividing by the number of genes in the clade. We compared both CBL and cumulative patristic distance between stable and unstable clades using a *t*-test. We tested for correlation between both CBL and cumulative patristic distance and the instability of a clade using Spearman’s rank correlation.

### Selection analyses

We split the dataset into the previously identified clades and carried out analysis individually on each clade. To construct a codon alignment for selection analyses we first aligned amino acid sequences using MAFFT v7.453 (-maxiterate 1000, using L-INS-I algorithm) (Katoh et al. 2002, 2005; Katoh & Standley 2013). We then used PAL2NAL to align the corresponding nucleotide sequences to the amino acid alignment (Suyama et al. 2006). Phylogenetic trees were constructed using IQ-TREE (Nguyen et al. 2015; Kalyaanamoorthy et al. 2017; Hoang et al. 2018).

### Gene conversion

As gene conversion can lead to false positive results when testing for positive selection with PAML, we first tested our dataset for evidence of gene conversion (Casola & Hahn 2009). We split the dataset by species and aligned the sequences from each species separately using MAFFT v7.453 (-maxiterate 1000, using L-INS-I algorithm) (Katoh et al. 2002, 2005; Katoh & Standley 2013). We tested for gene conversion using GENECONV in RDP5 (Martin et al. 2021; Padidam et al. 1999). We tested the sequences as linear sequences and found no evidence for gene conversion in our dataset.

#### Branch-specific models

To compare selection pressure on clades which have undergone expansions with those which have not we used codon substitution models implemented in Phylogenetic Analysis by Maximum Likelihood (PAML) (Yang 2007). Firstly, we carried out “one-ratio” models. For these, we assumed that dN/dS (the ω ratio of nonsynonymous to synonymous substitutions) has one value across the whole clade. We compared the dN/dS value between stable and unstable clades using a *t*-test, and tested for a correlation between the dN/dS value of a clade and its normalized CBL using Spearman’s rank correlation.

We then carried out “free-ratio” models, in which we allowed ω to vary between lineages. We compared the two models (“one-ratio” and “free-ratio) for each clade using a likelihood ratio test (LRT). To correct for multiple-testing, we used the p.adjust function in R, with false detection rate (fdr) correction, which controls for the proportion of false positives. To test if the unstable clades were more likely to be explained by a “free-ratio” model then stable clades we used a Fisher’s exact test.

Furthermore, to compare the selection pressures on individual genes or groups of genes within the clades, we again compared two models. The first assumed a single ω for the clade (“one-ratio”, as above), the second assumes two ω ratios, one for the lineage of interest, or group of interest, and the second for the rest of the tree. Again, we compared the two models using a likelihood ratio test (LRT), correcting for multiple-testing. To automate the process of testing each branch we used ETE3 framework to implement codeml (--leaves— internals –codeml_param CodonFreq,3 Nssites,0 fix_omega,0 omega,1 fix_kappa,0 kappa,0 cleandata,1 fix_blength,1) (Huerta-Cepas et al. 2016).

#### Site-specific models

To test for positively selected sites, we carried out two model comparisons: M1a vs. M2a, and M7 vs. M8. Model M1a is a nearly neutral model with two classes of sites, one evolving neutrally and the other under purifying selection. M2a is the same as M1a but with an additional site class for positive selection. Model M7 allows for a beta distribution of dN/dS across sites, while M8 has a beta distribution plus an additional site class with dN/dS>1 for positive selection. As above, we compared the two sets of models using likelihood ratio tests (LRT) and corrected for multiple-testing. We identified sites under positive selection using the Bayes Empirical Bayesian (BEB) in the PAML output (Yang et al. 2005). We considered sites to be under positive selection when the BEB posterior probability was greater than 0.9.

To determine where on the proteins these positively selected residues are located, we created protein models using ColabFold (Mirdita et al. 2021). ColabFold predicts protein structure based on AlphaFold2 (Jumper et al. 2021). We used the first sequence from each clade as a representative. We then visualized the resulting models using PyMOL (Schrödinger 2021).

We made figures using the packages *ggplot2* and *cowplot* in R version 3.6.2 (Wickham 2009, 2; R Core Team 2019).

## Data Availability Statement

Data and R scripts used for analysis are available from Open Science Framework (https://osf.io/9tdqu/?view_only=f7514c174fdb4574b77ed37ac1983f24).

## Acknowledgements

Santiago R. Ramírez was funded by the David and Lucile Packard Foundation. We thank members of the Ramírez lab for their feedback. We also thank Philipp Brand for his helpful comments and advice. We are grateful to Dave Curran who added the option to merge singletons to MiPhy at our request, and also Fabio Henrique Kuriki Mendes who helped with the interpretation of the CAFE results. We thank Lucas Rubio and Amelia Bassiti for allowing us to use their photos of *Melipona quadrifasciata* and *Osmia bicornia*, respectively.

## Notes

https://osf.io/9tdqu/?view_only=f7514c174fdb4574b77ed37ac1983f24

### Competing Interest Statement

The authors have declared no competing interest.

