## Supplementary information for "The birth-and-death evolution of cytochrome P450 genes in bees"

### Resumen en español

El modelo de nacimiento y muerte de la evolución de familias multigénicas describe cómo las familias pueden expandirse por duplicación y contraerse, tanto por perdida de genes como por formación de pseudogenes. Se cree que la estabilidad filogenética de un gen está relacionada con el grado de su importancia funcional. Sin embargo, no se sabe que tanto contribuyen los procesos adaptativos y neutrales a la evolución de los genes de familias multigénicas. Los citocromos P450 son una de las familias de genes más diversas y más ampliamente estudiadas. Son muy importantes para la desintoxicación y para otras funciones fisiológicas. Las abejas son un grupo de animales con elevada exposición a toxinas debido a su dieta rica en polen y néctar, en adición, algunas recolectan resinas. En este estudio, describimos los P450 de la abeja orquídea *Euglossa dilemma*. Las abejas orquídeas son un clado neotropical en el que los machos recolectan compuestos químicos del medio ambiente y los usan como feromonas, lo que resulta en una alta exposición a compuestos químicos. Realizamos análisis filogenéticos con diez especies de abejas de tres familias de abejas. No encontramos relación entre el repertorio P450 de una abeja y su ecología. Además, el análisis revela que los clados P450 pueden clasificarse en clados estables e inestables, y que es más probable que los genes implicados en el metabolismo xenobiótico pertenezcan a clados inestables. Además, encontramos que los clados inestables están bajo presiones evolutivas más dinámicas, con señales de evolución adaptativa detectadas. Esto sugiere que tanto la duplicación de genes como la divergencia de secuencia bajo selección positiva han desempeñado un papel en la diversificación de las P450. Nuestro trabajo resalta que la complejidad de la evolución familiar multigénica no siempre sigue predicciones generalizadas.

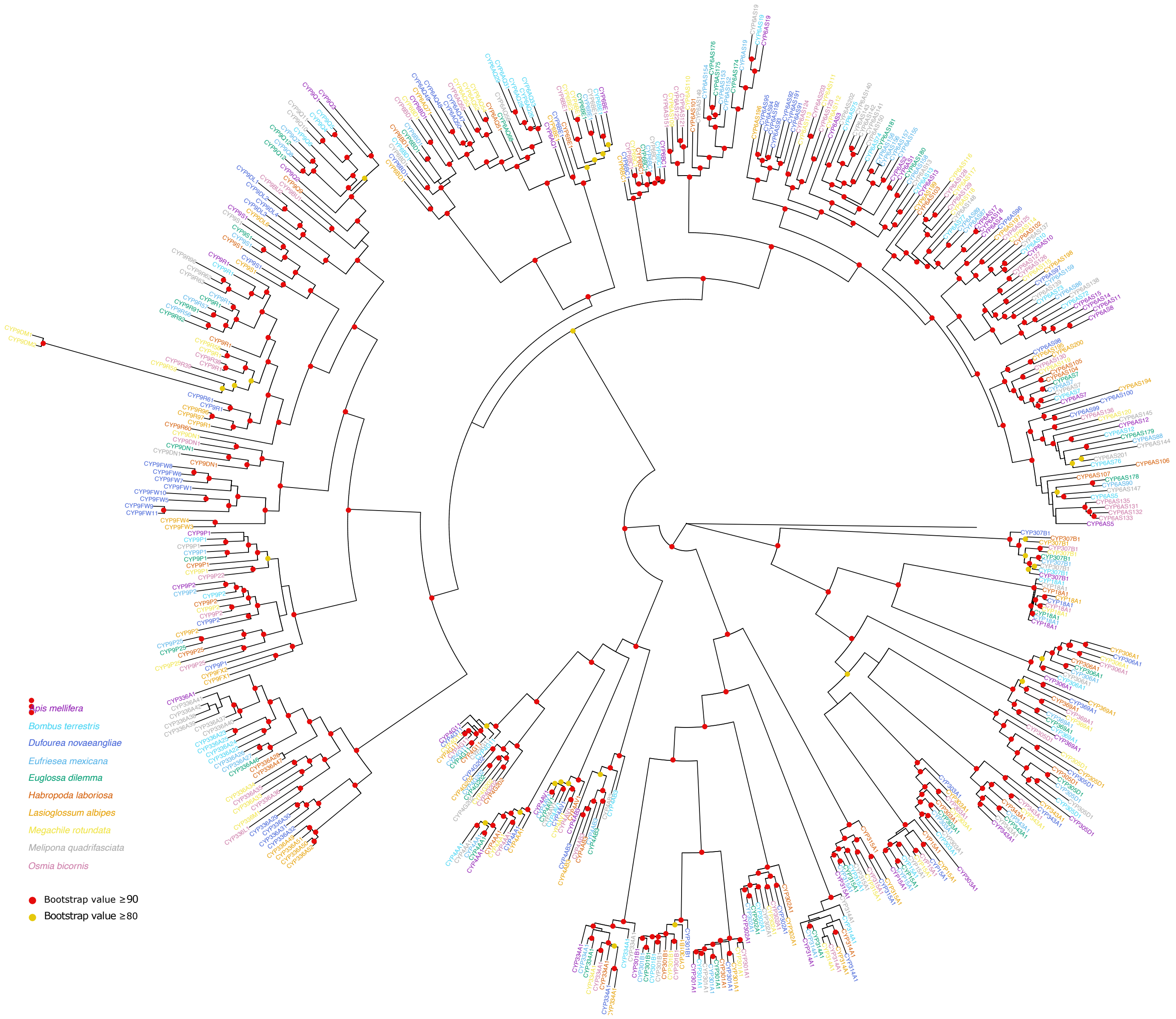


***Supplementary Figure 1:*** *Full phylogeny of cytochrome P450s in bees. The phylogeny was constructed in IQ -TREE (model JTT+F+R10) using 481 amino acid sequences across ten species of bee. Bootstrap values (n=1000) are illustrated.*

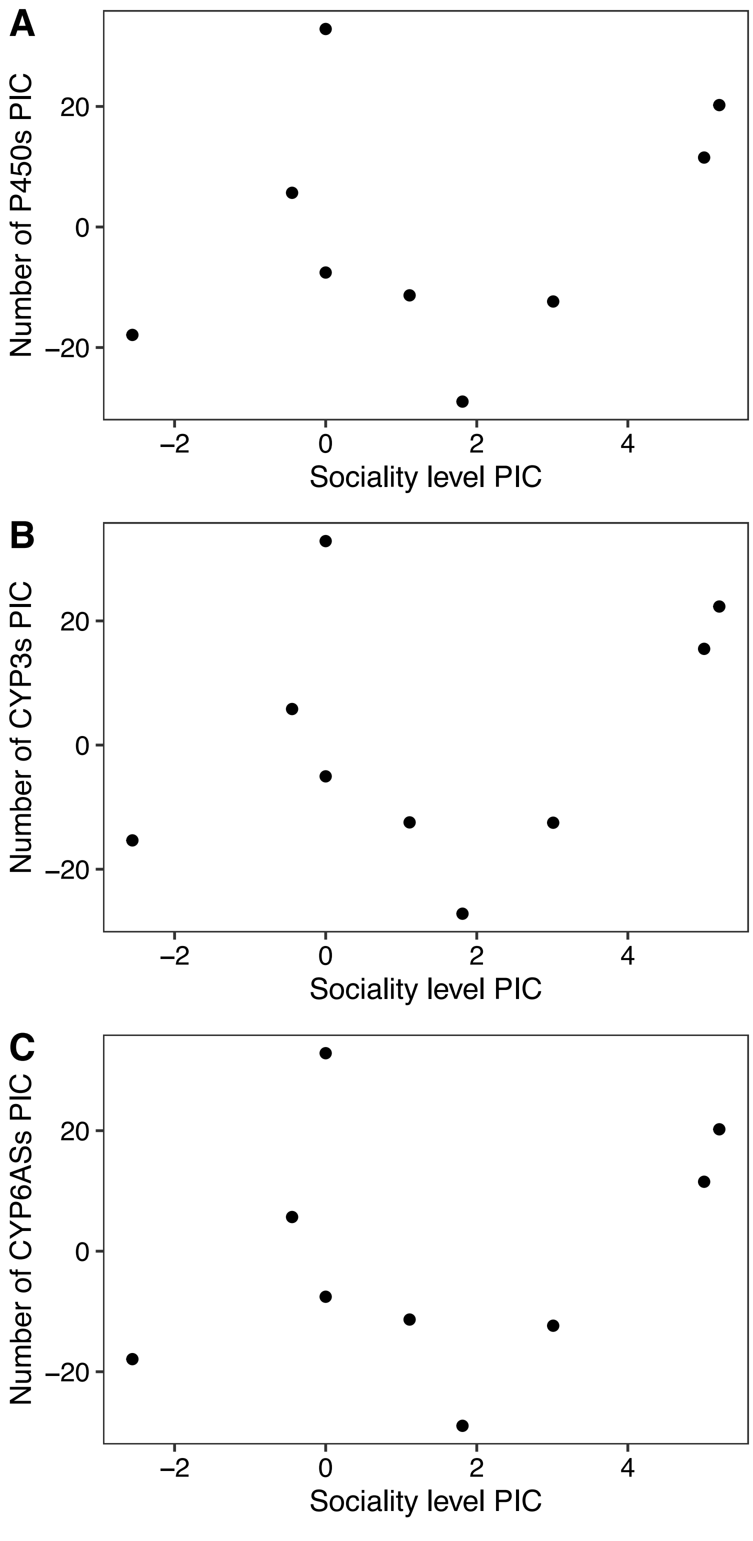


***Supplementary Figure 2:*** *Relationship between sociality and P450 repertoire. We did not find a relationship between sociality and A) the number of P450s (Spearman's rank correlation, S=93.891, p=0.5739, rho=0.218), B) the number of CYP3s (Spearman's rank correlation, S=93.891, p=0.5739, rho=0.218), or C) the number of CYP6AS genes (Spearman's rank correlation, S=52.719, p=0.1163, rho=0.5607). PIC, phylogenetically independent contrast.*

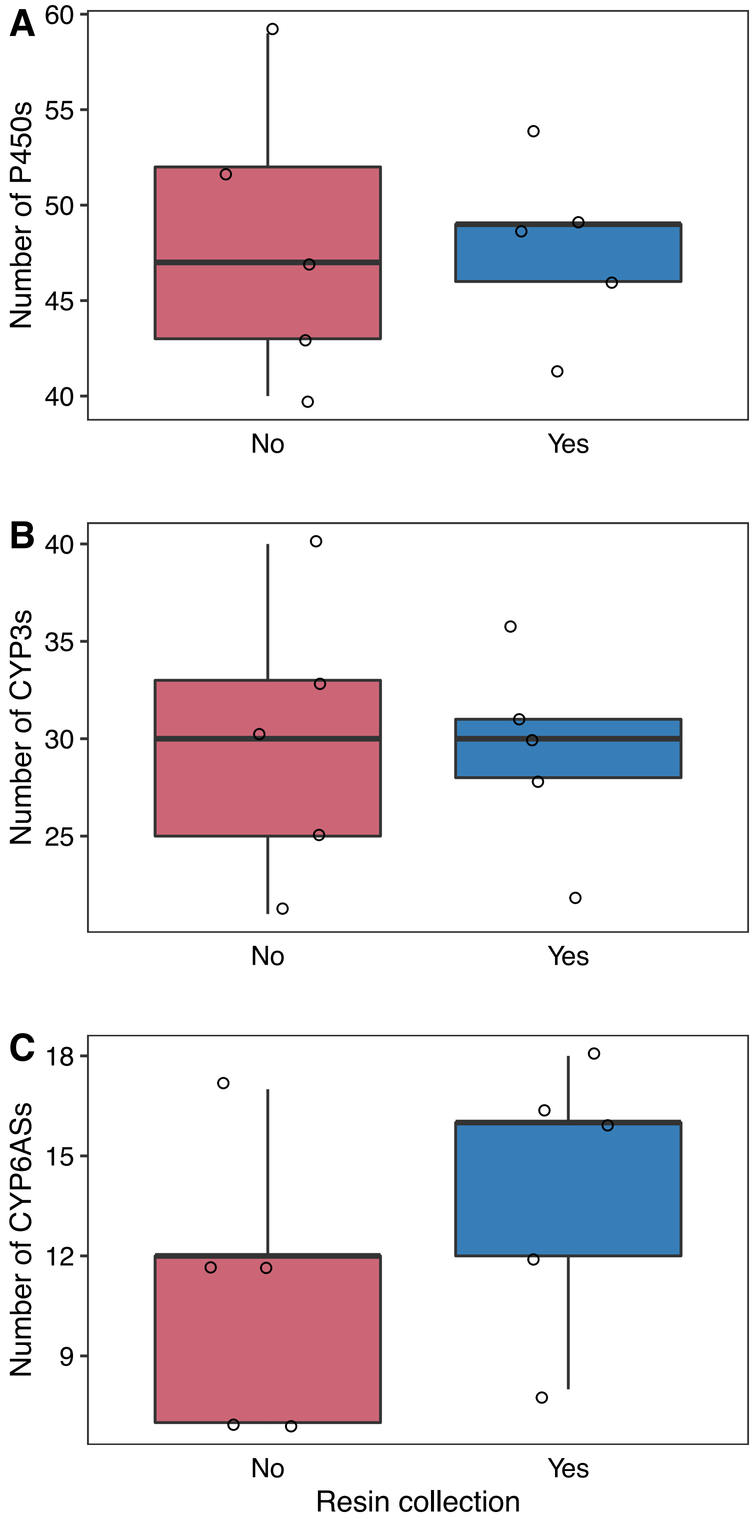


***Supplementary Figure 3:*** *Boxplots showing relationship between resin collection and P450 repertoire. We did not find a relationship between resin collection and A) the number of P450s (phylogenetic ANOVA, F=0.01008, df=1, p=0.94), B) the number of CYP3s (phylogenetic ANOVA, F=0.01006, df=1, p=0.94), or C) the number of CYP6AS genes (phylogenetic ANOVA, F=1.3433, df=1, p=0.366). PIC, phylogenetically independent contrast.*

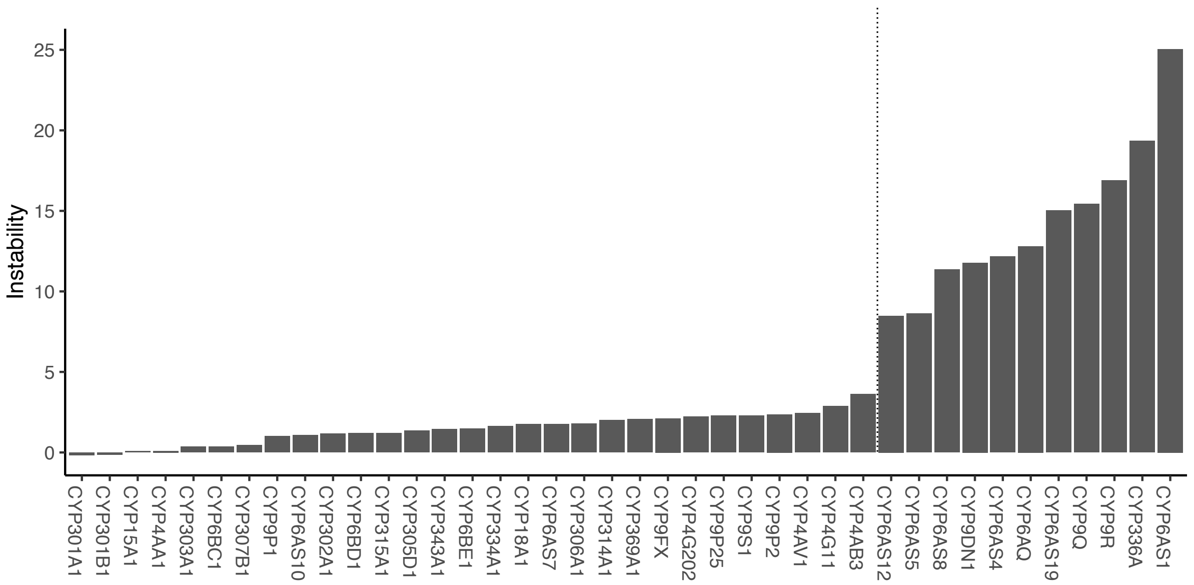


***Supplementary Figure 4:*** *Distribution of instability scores across P450 clades. Instability scores from MiPhy output plotted for each clade. Clades to the left of the dotted line are considered stable, and those to the right unstable.*

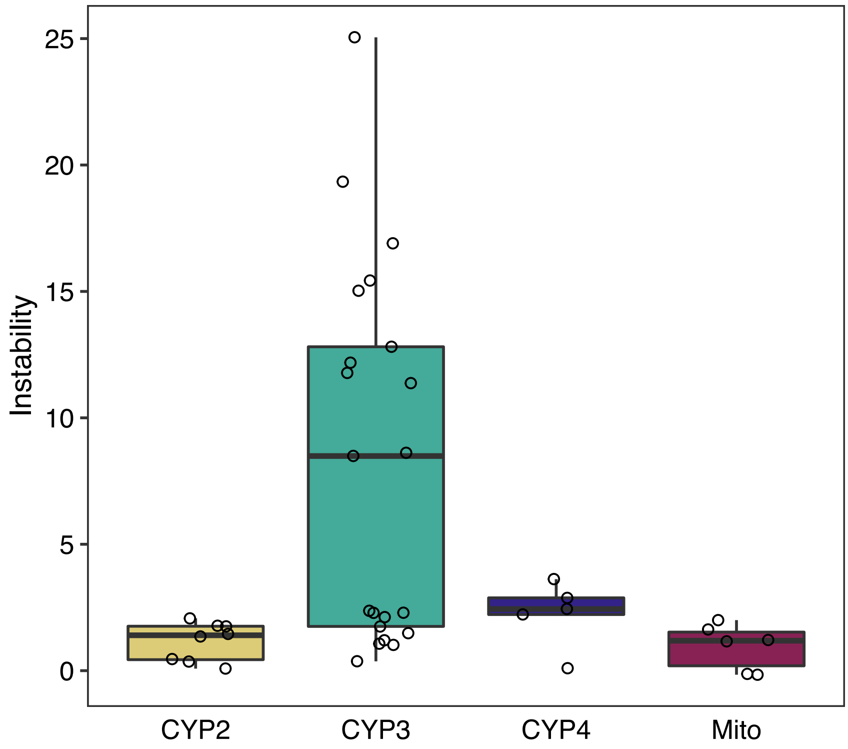


***Supplementary Figure 5:*** *Boxplot showing MiPhy instability scores across the four CYP groups. Instability differs between the four groups (Welch one-way ANOVA test, $F_{3,13.6}$=6.8, p=0.005). A Games-Howell post-hoc analysis revealed statistically significant increases in instability in the CYP3 group when compared with the other three groups CYP2 (7.07, 95% CI 2.54-11.6, p=*0.001*), CYP4 (5.98, 95% CI 1.26-10.7, p=0.009), and mitochondrial (7.28, 95% CI 2.71-11.9, 0.001). Mito; mitochondrial.*

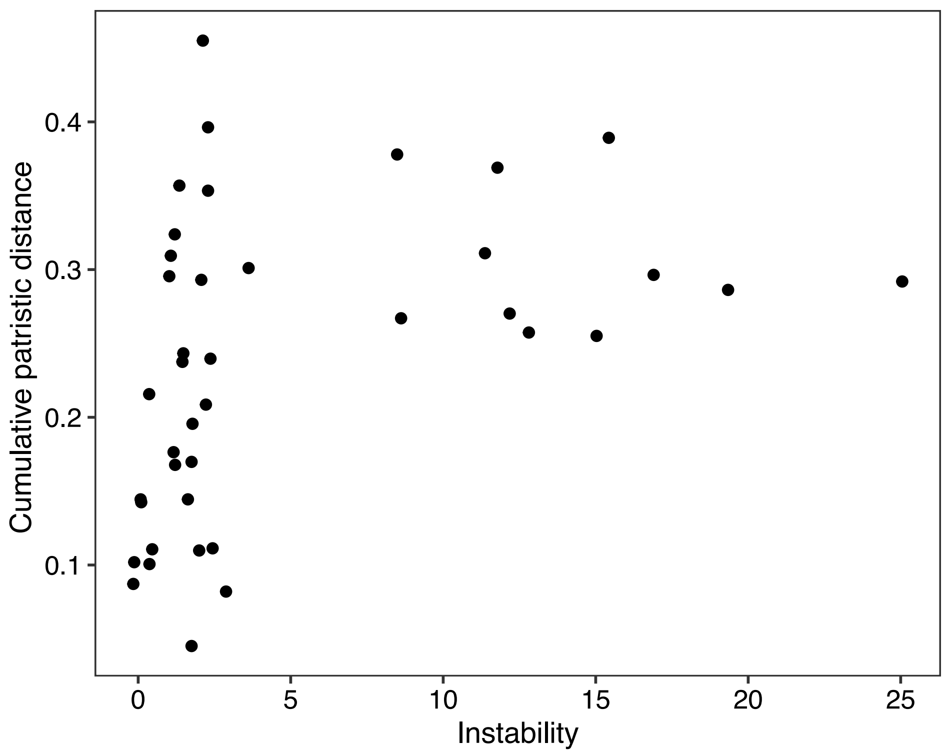


***Supplementary Figure 6:*** *Relationship between cumulative patristic distance and the instability of a clade. When cumulative patristic distance is normalized for gene number in a clade, there is a significant correlation between clade cumulative patristic distance and clade instability (Spearman's Correlation, rho=0.475, p= 0.00191). CBL, cumulative branch length.*

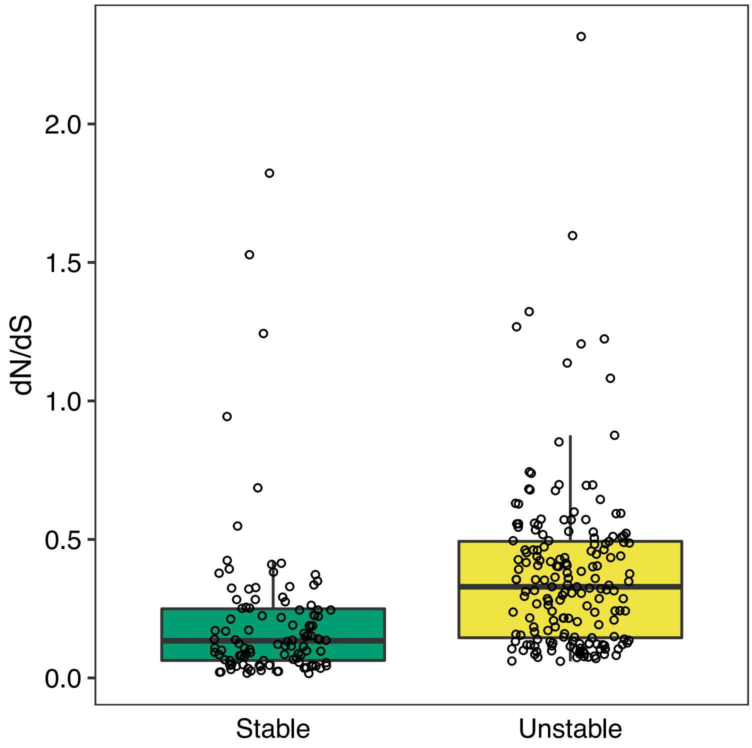


***Supplementary Figure 7:*** *Boxplot showing distribution of all branches with dN/dS significantly different from background dN/dS of the clade (adjusted p value<0.05). Branches with a dN/dS>10 were removed. The dN/dS of branches found in unstable clades is significantly higher than that of stable clades (*t*-test, df=238.2, t=-4.837, p=0.000002364).*

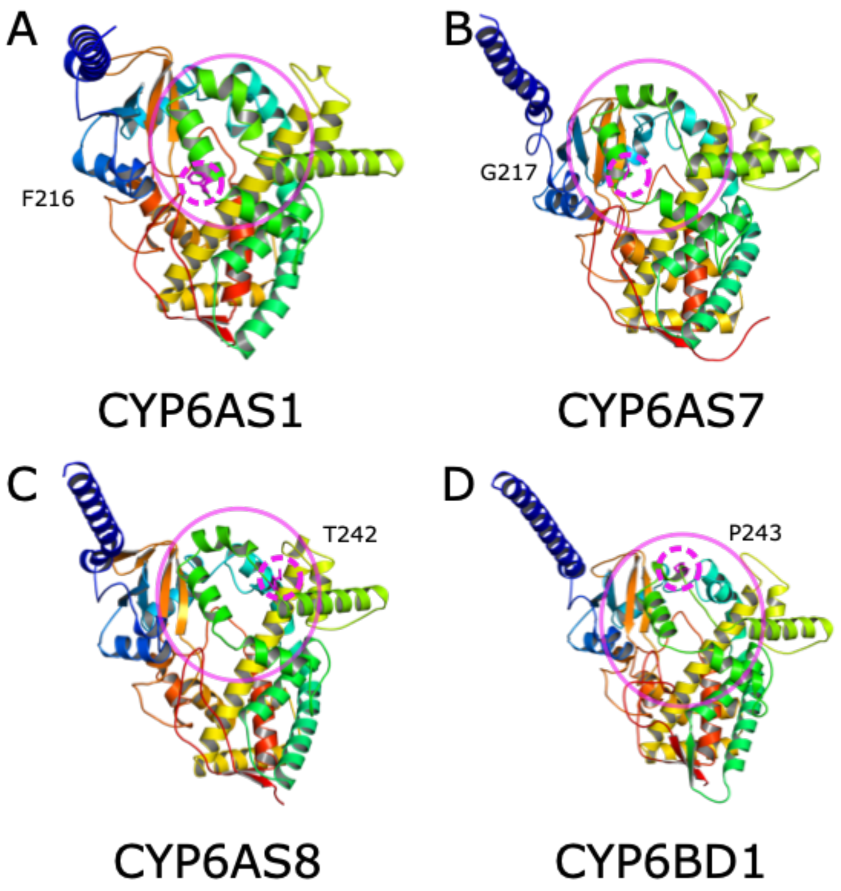


***Supplementary Figure 8:*** *Protein models for representative P450s for the clades where positive selection was detected in PAML site models. Structures shown for A) CYP6AS1 (GB40287), B) CYP6AS7 (GB49894), C) CYP6AS8 (GB49878) and D) CYP6BD1 (GB47279). Positively selected residues are highlighted by the pink dashed line with the surface area in which all residues are found highlighted with the solid line circle.*

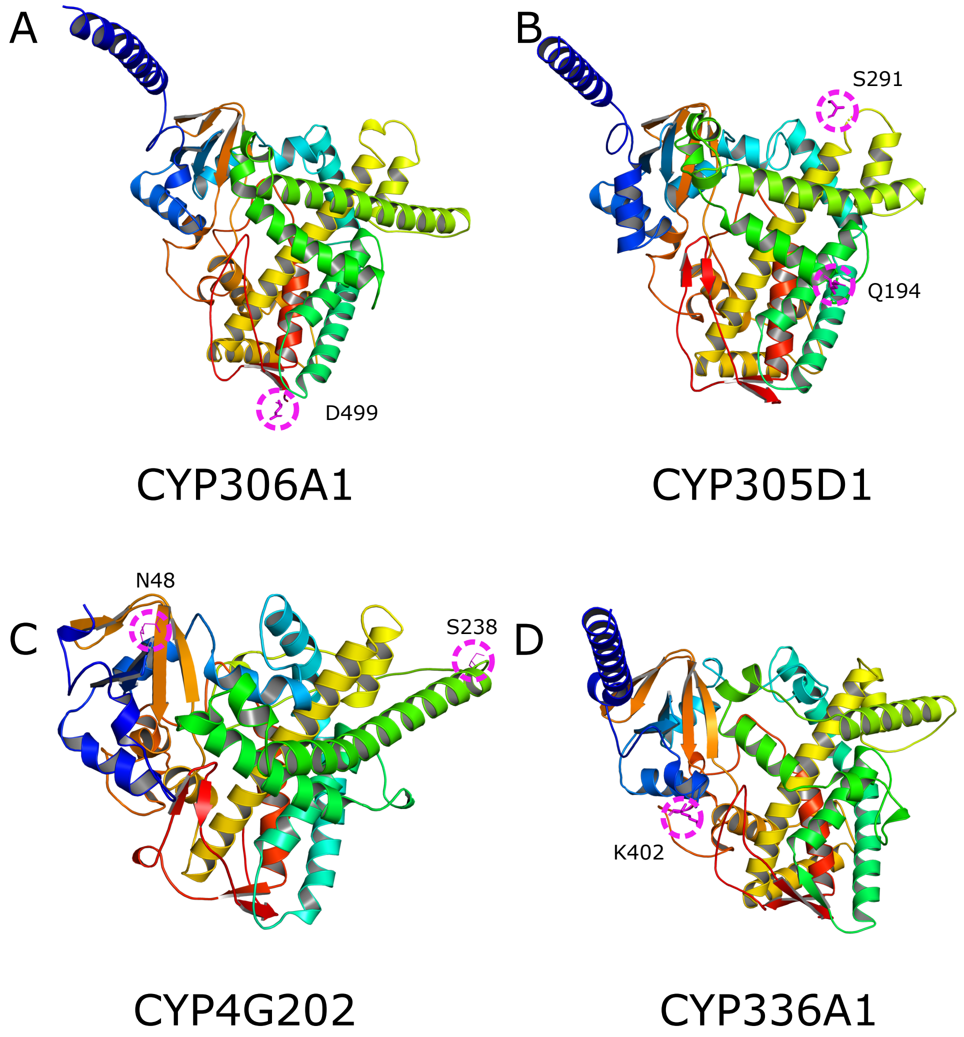


***Supplementary Figure 9:*** *Protein models for representative P450s for the clades where positive selection was detected in PAML site models. Structures shown for A) CYP306A1 (GB54743), B) CYP305D1 (GB48105), C) CYP4G202 (XM_015579213.1) and D) CYP336A1 (GB55669). Positively selected residues are highlighted by the pink dashed line.*

***Supplementary Table 1***: *Results from CAFE analysis. Only clades with significant posterior probabilities are shown, full results available from OSF (https://osf.io/9tdqu/?view_only=f7514c174fdb4574b77ed37ac1983f24).*

| #FamilyID | Gamma Cat Median | Likelihood of Category | Likelihood of Family | Posterior Probability | Significant |
|---|---|---|---|---|---|
| CYP336A | 0.0170496 | 3.98E-24 | 5.92E-11 | 6.72E-14 | N/S |
| CYP336A | 0.311338 | 3.07E-14 | 5.92E-11 | 0.00051822 | N/S |
| CYP336A | 2.67161 | 5.92E-11 | 5.92E-11 | 0.999482 | * |
| CYP9DN1 | 0.0170496 | 5.24E-28 | 4.81E-10 | 1.09E-18 | N/S |
| CYP9DN1 | 0.311338 | 6.71E-16 | 4.81E-10 | 1.40E-06 | N/S |
| CYP9DN1 | 2.67161 | 4.81E-10 | 4.81E-10 | 0.999999 | * |
| CYP9R | 0.0170496 | 5.59E-21 | 7.46E-10 | 7.50E-12 | N/S |
| CYP9R | 0.311338 | 6.91E-12 | 7.46E-10 | 0.00926094 | N/S |
| CYP9R | 2.67161 | 7.39E-10 | 7.46E-10 | 0.990739 | * |
| CYP9Q | 0.0170496 | 1.15E-21 | 1.55E-09 | 7.45E-13 | N/S |
| CYP9Q | 0.311338 | 2.46E-12 | 1.55E-09 | 0.0015898 | N/S |
| CYP9Q | 2.67161 | 1.54E-09 | 1.55E-09 | 0.99841 | * |
| CYP6AQ | 0.0170496 | 3.91E-29 | 8.73E-11 | 4.47E-19 | N/S |
| CYP6AQ | 0.311338 | 5.27E-17 | 8.73E-11 | 6.03E-07 | N/S |
| CYP6AQ | 2.67161 | 8.73E-11 | 8.73E-11 | 0.999999 | * |
| CYP6AS1 | 0.0170496 | 2.79E-42 | 3.85E-13 | 7.25E-30 | N/S |
| CYP6AS1 | 0.311338 | 4.68E-22 | 3.85E-13 | 1.21E-09 | N/S |
| CYP6AS1 | 2.67161 | 3.85E-13 | 3.85E-13 | 1 | * |
| CYP6AS8 | 0.0170496 | 1.18E-21 | 5.51E-10 | 2.14E-12 | N/S |
| CYP6AS8 | 0.311338 | 2.63E-13 | 5.51E-10 | 0.00047735 | N/S |
| CYP6AS8 | 2.67161 | 5.51E-10 | 5.51E-10 | 0.999523 | * |
| CYP6AS4 | 0.0170496 | 1.92E-21 | 1.44E-09 | 1.33E-12 | N/S |
| CYP6AS4 | 0.311338 | 4.53E-13 | 1.44E-09 | 0.000315 | N/S |
| CYP6AS4 | 2.67161 | 1.44E-09 | 1.44E-09 | 0.999685 | * |

***Supplementary Table 2**: Results from codeml analysis of all clades comparing "one-ratio" and "free-ratio" models. Log likelihood (lnL1 for "one-ratio", lnL2 for "free-ratio"), as well as number of parameters (np1 for "one-ratio", np2 for "free-ratio") are shown for both models. Twice the difference in log likelihood score (2*ΔlnL) is used along with the difference in number of parameters (Δnp) as the degrees of freedom to carry out a chi-squared test to compare the models. The adjusted p-value (adj. p-val) is corrected using false-detection rate correction.*

| Clade | Type | lnL1 | np1 | lnL2 | np2 | 2*ΔLnl | Δnp | adj. p-val |
|---|---|---|---|---|---|---|---|---|
| 1 | Stable | -8885.521119 | | -8877.035735 | | 16.970904 | 16 | NS |
| 2 | Stable | -7595.834319 | | -7576.617635 | | 38.43349 | 16 | 0.00194 |
| 3 | Stable | -4276.055519 | | -4255.898535 | | 40.31409 | 16 | 0.00117 |
| 4 | Stable | -7277.450619 | | -7256.954935 | | 40.991366 | 16 | 0.000969 |
| 5 | Stable | -5443.280819 | | -5433.125835 | | 20.310038 | 16 | NS |
| 6 | Stable | -8222.549119 | | -8212.113335 | | 20.871602 | 16 | NS |
| 7 | Stable | -6158.749219 | | -6146.421135 | | 24.656106 | 16 | NS |
| 8 | Stable | -5894.655719 | | -5882.141835 | | 25.027812 | 16 | NS |
| 9 | Stable | -8893.052519 | | -8875.586335 | | 34.932424 | 16 | 0.00524 |
| 10 | Stable | -9143.509917 | | -9118.719 | 31 | 49.581888 | 14 | 1.43E-05 |
| 11 | Stable | -9995.690819 | | -9976.173535 | | 39.034538 | 16 | 0.00165 |
| 12 | Stable | -5551.574717 | | -5530.669631 | | 41.810198 | 14 | 0.000241 |
| 13 | Stable | -6872.413119 | | -6824.283135 | | 96.260048 | 16 | 5.33E-13 |
| 14 | Stable | -7565.741819 | | -7548.038 | 35 | 35.407436 | 16 | 0.00466 |
| 15 | Stable | -9248.604917 | | -9200.806731 | | 95.59643 | 14 | 1.20E-13 |
| 16 | Stable | -7084.257417 | | -7066.173431 | | 36.16804 | 14 | 0.00158 |
| 17 | Stable | -8112.432719 | | -8066.574635 | | 91.71616 | 16 | 3.22E-12 |
| 18 | Stable | -7385.827715 | | -7357.991227 | | 55.672872 | 12 | 3.22E-07 |
| 19 | Stable | -8588.996 | 19 | -8539.473335 | | 99.045384 | 16 | 1.74E-13 |
| 20 | Unstable | -13023.38 | 59 | -12939.732115 | | 167.2961 | 56 | 1.46E-12 |
| 21 | Stable | -4553.68415 | | -4553.233 | 7 | 0.902394 | 2 | NS |
| 22 | Stable | -6548.36339 | | -6541.832915 | | 13.060756 | 6 | NS |
| 23 | Stable | -8640.813815 | | -8599.545327 | | 82.537118 | 12 | 3.38E-12 |
| 24 | Stable | -6546.013915 | | -6531.013527 | | 30.000666 | 12 | 0.00385 |
| 25 | Unstable | -15955.60529 | | -15842.42855 | | 226.35529226 | | 7.25E-33 |
| 26 | Stable | -8377.550613 | | -8368.515823 | | 18.069706 | 10 | NS |
| 27 | Unstable | -21124.72251 | | -20935.45599 | | 378.53361 | 48 | 2.63E-51 |
| 28 | Unstable | -20864.27439 | | -20736.59675 | | 255.35556436 | | 9.76E-34 |
| 29 | Stable | -10640.31419 | | -10609.68435 | | 61.260314 | 16 | 6.75E-07 |
| 30 | Unstable | -16679.89135 | | -16552.37667 | | 255.02871 | 32 | 2.77E-35 |
| 31 | Stable | -8462.225421 | | -8442.611939 | | 39.226992 | 18 | 0.00379 |
| 32 | Stable | -7188.234919 | | -7163.877935 | | 48.714076 | 16 | 6.98E-05 |
| 33 | Unstable | -14376.64733 | | -14303.31963 | | 146.65717430 | | 1.34E-16 |
| 34 | Unstable | -17683.19269 | | -17560.453135 | | 245.47670666 | | 1.19E-21 |
| 35 | Stable | -8131.841815 | | -8125.223127 | | 13.237244 | 12 | NS |
| 36 | Unstable | -15279.11929 | | -15214.61 | 55 | 129.01923 | 26 | 5.11E-15 |
| 37 | Unstable | -11734.96427 | | -11697.21151 | | 75.505834 | 24 | 6.75E-07 |
| 38 | Stable | -9591.545723 | | -9512.079343 | | 158.93278420 | | 9.65E-23 |
| 39 | Unstable | -10903.74925 | | -10840.50547 | | 126.48877 | 22 | 5.07E-16 |
| 40 | Unstable | -9711.623 | 21 | -9651.110639 | | 121.02476618 | | 1.34E-16 |

***Supplementary Table 3***: *Results from codeml analysis of all clades comparing M7 and M8 models to test for positively selected sites. Log likelihood (lnL1 for "one-ratio", lnL2 for "free-ratio"), as well as number of parameters (np1 for M7, np2 for M8 are shown for both models. Twice the difference in log likelihood score (2*ΔlnL) is used along with the difference in number of parameters (Δnp) as the degrees of freedom to carry out a chi-squared test to compare the models. The adjusted p-value (adj. p-val) is corrected using false-detection rate correction.*

| Clade | Type | lnL1 | np1 | lnL2 | np2 | 2*ΔLnl | Δnp | adj. p-val |
|---|---|---|---|---|---|---|---|---|
| 1 | Stable | -8766.659 | 20 | -8766.6603 | 22 | -0.002622 | 2 | NS |
| 2 | Stable | -7455.4438 | 20 | -7453.4867 | 22 | 3.914212 | 2 | NS |
| 3 | Stable | -4141.5873 | 20 | -4139.7375 | 22 | 3.699756 | 2 | NS |
| 4 | Stable | -7174.9928 | 20 | -7172.9778 | 22 | 4.03005 | 2 | NS |
| 5 | Stable | -5357.2036 | 20 | -5356.6783 | 22 | 1.050562 | 2 | NS |
| 6 | Stable | -8071.0699 | 20 | -8071.0017 | 22 | 0.136368 | 2 | NS |
| 7 | Stable | -6022.6043 | 20 | -6022.6063 | 22 | -0.003978 | 2 | NS |
| 8 | Stable | -5835.0172 | 20 | -5832.4644 | 22 | 5.105696 | 2 | NS |
| 9 | Stable | -8636.7597 | 20 | -8631.9576 | 22 | 9.604192 | 2 | 0.0365 |
| 10 | Stable | -8981.8821 | 18 | -8980.4673 | 20 | 2.829626 | 2 | NS |
| 11 | Stable | -9812.4871 | 20 | -9807.6264 | 22 | 9.721406 | 2 | 0.0365 |
| 12 | Stable | -5468.6527 | 18 | -5465.1472 | 20 | 7.011098 | 2 | NS |
| 13 | Stable | -6757.9741 | 20 | -6754.8488 | 22 | 6.250688 | 2 | NS |
| 14 | Stable | -7468.6328 | 20 | -7468.6333 | 22 | -0.001046 | 2 | NS |
| 15 | Stable | -8992.6945 | 18 | -8991.1039 | 20 | 3.181284 | 2 | NS |
| 16 | Stable | -7006.6508 | 18 | -7006.544 | 20 | 0.213514 | 2 | NS |
| 17 | Stable | -7944.9224 | 20 | -7935.4476 | 22 | 18.94961 | 2 | 0.001536 |
| 18 | Stable | -7169.0795 | 16 | -7164.4796 | 18 | 9.199732 | 2 | 0.0402128 |
| 19 | Stable | -8422.2002 | 20 | -8419.699 | 22 | 5.002442 | 2 | NS |
| 20 | Unstable | -12609.596 | 60 | -12600.891 | 62 | 17.409332 | 2 | 0.00221067 |
| 21 | Stable | -4518.7153 | 6 | -4516.9636 | 8 | 3.5034 | 2 | NS |
| 22 | Stable | -6468.1836 | 10 | -6467.3351 | 12 | 1.696942 | 2 | NS |
| 23 | Stable | -8495.2221 | 16 | -8493.3291 | 18 | 3.785992 | 2 | NS |
| 24 | Stable | -6405.2683 | 16 | -6405.2688 | 18 | -0.00098 | 2 | NS |
| 25 | Unstable | -15647.485 | 30 | -15647.334 | 32 | 0.300714 | 2 | NS |
| 26 | Stable | -8272.2157 | 14 | -8272.0471 | 16 | 0.337172 | 2 | NS |
| 27 | Unstable | -20591.579 | 52 | -20585.675 | 54 | 11.809022 | 2 | 0.01558343 |
| 28 | Unstable | -20345.655 | 40 | -20345.581 | 42 | 0.148884 | 2 | NS |
| 29 | Stable | -10428.48 | 20 | -10420.618 | 22 | 15.722438 | 2 | 0.003854 |
| 30 | Unstable | -16350.332 | 36 | -16347.063 | 38 | 6.537426 | 2 | NS |
| 31 | Stable | -8302.8489 | 22 | -8300.7518 | 24 | 4.194036 | 2 | NS |
| 32 | Stable | -7105.6278 | 20 | -7105.6303 | 22 | -0.005126 | 2 | NS |
| 33 | Unstable | -14123.312 | 34 | -14122.254 | 36 | 2.11482 | 2 | NS |
| 34 | Unstable | -17127.962 | 70 | -17117.188 | 72 | 21.548134 | 2 | 0.000836 |
| 35 | Stable | -7977.2978 | 16 | -7977.2627 | 18 | 0.070212 | 2 | NS |
| 36 | Unstable | -14949.979 | 30 | -14943.968 | 32 | 12.021974 | 2 | 0.01558343 |
| 37 | Unstable | -11406.922 | 28 | -11404.484 | 30 | 4.87628 | 2 | NS |
| 38 | Stable | -9417.3754 | 24 | -9410.5437 | 26 | 13.663456 | 2 | 0.008632 |
| 39 | Unstable | -10687.727 | 26 | -10686.097 | 28 | 3.259078 | 2 | NS |
| 40 | Unstable | -9555.8937 | 22 | -9552.2736 | 24 | 7.240104 | 2 | NS |
